# Type II Fusarium head blight susceptibility factor identified in wheat

**DOI:** 10.1101/2020.02.06.937425

**Authors:** B. Hales, A. Steed, V. Giovannelli, C. Burt, M. Lemmens, M. Molnár-Láng, P. Nicholson

## Abstract

Fusarium head blight (FHB) causes significant grain yield and quality reductions in wheat and barley. Most wheat varieties are incapable of preventing FHB spread through the rachis, but disease is typically limited to individually infected spikelets in barley. We point inoculated wheat lines possessing barley chromosome introgressions to test whether FHB resistance could be observed in a wheat genetic background. The most striking differential was between 4H(4D) substitution and 4H addition lines. The 4H addition line was similarly susceptible to the wheat parent, but the 4H(4D) substitution line was highly resistant, which suggests that there is an FHB susceptibility factor on wheat chromosome 4D. Point inoculation of Chinese Spring 4D ditelosomic lines demonstrated that removing 4DS results in high FHB resistance. We genotyped four Chinese Spring 4DS terminal deletion lines to better characterise the deletions in each line. FHB phenotyping indicated that lines del4DS-2 and del4DS-4, containing smaller deletions, were susceptible and had retained the susceptibility factor. Lines del4DS-3 and del4DS-1 contain larger deletions and were both significantly more resistant, and hence had presumably lost the susceptibility factor. Combining the genotyping and phenotyping results allowed us to refine the susceptibility factor to a 31.7 Mbp interval on 4DS.

**Highlight:** We have identified a Type II Fusarium head blight susceptibility factor on the short arm of wheat chromosome 4D and refined its position to a 31.7 Mbp interval.

## Introduction

Fusarium head blight (FHB) is an economically important fungal disease of various cereal crop species, in particular wheat (*Triticum aestivum*) and barley (*Hordeum vulgare).* In wheat, the primary symptom is the premature bleaching of spikelets that progressively spreads through the head. Infected spikelets produce shrivelled and chalky grain, which can have a significant impact on yield. Furthermore, mycotoxins accumulate in infected grain, which are harmful to humans and animal consumers. The most important mycotoxin is deoxynivalenol (DON) which acts as a virulence factor in wheat by promoting the spread of the fungus (Bai *et al.*, 2002; Langevin *et al.*, 2004). *Fusarium graminearum* and *F. culmorum* are the most prevalent species responsible for FHB. Both species are capable of producing large quantities of DON (Scherm *et al.*, 2013) and hence tend to be the most aggressive pathogens of wheat.

Resistance to initial infection (Type I) and to the spread of infection through the rachis (Type II) were first proposed by Schroeder and Christensen (1963) and remain the two most widely considered forms of resistance. Numerous small-effect Type II and fewer Type I FHB quantitative trait loci (QTL) have been reported and are reviewed by Buerstmayr *et al.* (2009) and more recently by Buerstmayr *et al.* (2019). In addition to these two main types of FHB resistance, there is resistance to kernel infection (Type III), host tolerance to FHB and/ or DON (Type IV) and resistance to the accumulation of DON (Type V) (Boutigny *et al.*, 2008; Gunupuru *et al.*, 2017). Single amino acid changes to the DON target, ribosomal protein L3 (RPL3), have been demonstrated to improve tolerance to DON in yeast and hence this is a possible target to improve type IV resistance (Lucyshyn *et al.*, 2007; Mitterbauer *et al.*, 2004). Type V resistance is commonly considered to be a component of Type II resistance, as it typically limits disease spread (Gunupuru *et al.*, 2017), and can be subdivided into Class 1: processes that chemically modify DON to a less toxic form, and Class 2: processes that prevent the accumulation of DON and other trichothecene mycotoxins (Boutigny *et al.*, 2008). The most widely reported form of host detoxification of DON is by UDP-glucosyltransferase (UGT) proteins, which glucosylate DON to the less toxic DON-3-*O*-glucoside (D3G) (Poppenberger *et al.*, 2003). More recent studies have identified other pathways capable of detoxifying DON. For example, bacterial aldo-keto reductases were demonstrated to be involved in epimerising DON to 3-*epi*-DON (Hassan *et al.*, 2017; He *et al.*, 2017).

Wheat and barley differ noticeably in Type II resistance. Wheat typically possesses some degree of Type II susceptibility whilst, in contrast, barley is generally highly resistant to fungal spread through the rachis (Langevin *et al.*, 2004). Furthermore, whilst DON has been shown to function as a virulence factor in wheat (Langevin *et al.*, 2004), DON does not appear to possess such a role during infection of barley heads (Maier *et al.*, 2006).

The reasons for this marked difference in Type II susceptibility of wheat and barley are not well understood. Defined genetic stocks of wheat containing all or part of barley chromosomes offers an insight into which barley chromosomes contribute most strongly to Type II FHB resistance and whether this resistance can be expressed, and potentially utilised, in a wheat genetic background. Herein, we report on a series of experiments to establish whether this difference in FHB susceptibility is because barley carries genes conferring resistance, wheat carries genes conferring susceptibility, or whether it is a combination of both factors. Following this, we investigated the location of a major effect identified on wheat chromosome 4D that appears to significantly compromise resistance to disease spread through the rachis (Type II resistance).

To date, there have been few reports of FHB susceptibility factors. Garvin *et al.* (2015) identified a spontaneous deletion of a portion of the long arm of 3D, which appeared to be responsible for increased FHB resistance, suggesting that the deleted region carries an FHB susceptibility factor in the cultivar Apogee. Ma *et al.* (2006) point inoculated the existing ditelosomic lines of Chinese Spring that each lack individual chromosome arms. They found that the loss of individual chromosome arms can improve, as well as compromise, FHB resistance (Ma *et al.*, 2006). Their data suggested that some chromosome arms, especially 7AS, 3BL, 7BS and 4DS, are likely to contain FHB susceptibility factors (Ma *et al.*, 2006). Although the gene(s) underlying *Fhb1*, the most widely deployed FHB resistance QTL, remains controversial, there is evidence that *Fhb1* may be considered a disrupted susceptibility factor (Su *et al.*, 2019; Su *et al.*, 2018). Plant hormones play an important role in responding to disease. Host response to FHB infection is particularly sensitive to disrupting phytohormone production or perception. Plants insensitive to ethylene and brassinosteroid signalling exhibits increased FHB resistance, suggesting that the fungus is exploiting such physiological processes (Chen *et al.*, 2009; Goddard *et al.*, 2014). There is significant potential in identifying and characterising susceptibility factors, with the aim of eliminating them from elite cultivars to enhance resistance to FHB and other economically important diseases.

## Materials and Methods

### Plant material

Wheat-barley addition, substitution and translocation lines were developed at the Hungarian Academy of Sciences, Agricultural Institute, Centre for Agricultural Research, Hungary (Table 1). An independent set of wheat-barley addition lines, of the wheat variety Chinese Spring and the barley donor variety Betzes, were generated by Islam *et al.* (1981) and obtained from the Genetic Resources Unit at the John Innes Centre, Norwich, UK.

**Table 1.** Wheat-barley addition, substitution, translocation and centric fusion lines used in FHB experiments. The primary wheat parent was Martonvasari 9 kr1 (Mv9kr1) for all lines and the barley donor parents were Igri or Betzes. Associated references contain detailed descriptions of line generation and composition.

| Line abbreviation | Description | Reference |
| --- | --- | --- |
| Mv9kr1 | Martonvasari9 *kr1* | MolnarLang *et al.* (1996) |
| 1HS add | Mv9kr1–Igri 1HS disomic addition | Szakacs and Molnar-Lang (2007) |
| 2H add | Mv9kr1–Igri 2H disomic addition | Szakacs and Molnar-Lang (2007) |
| 3H add | Mv9kr1–Igri 3H disomic addition | Szakacs and Molnar-Lang (2007) |
| 4H add | Mv9kr1–Igri 4H disomic addition | Szakacs and Molnar-Lang (2007) |
| 6HS add | Mv9kr1–Igri 6HS disomic addition | Szakacs and Molnar-Lang (2010) |
| 7H add | Mv9kr1–Igri 7H disomic addition | Szakacs and Molnar-Lang (2010) |
| 2D-1H trans | 2DS.2DL-1HS translocation | Nagy *et al.* (2002) |
| 3HS.3BL centric | 3HS.3BL centric fusion | Nagy *et al.* (2002) |
| 4H(4D) sub | 4H(4D) wheat-barley substitution | Molnar *et al.* (2007) |
| 6B-4H trans | 6BS.6BL–4HL translocation | Nagy *et al.* (2002) |
| 7D-5H trans | 5HS-7DS.7DL wheat-barley translocation | Kruppa *et al.* (2013) |

Chinese Spring and its 4D ditelosomic (DT) lines were acquired from the Germplasm Resource Unit, John Innes Centre, Norwich, UK. The lines DT(4DL) and DT(4DS) lack 4DS and 4DL, respectively. Four homozygous Chinese Spring terminal deletion lines of 4DS, described by Endo and Gill (1996), were obtained from Kansas State University, USA. The lines acquired were 4532 L1 (FL= 0.53), 4532 L2 (FL= 0.82), 4532 L3 (FL= 0.67) and 4532 L4 (FL= 0.77), henceforth referred to as del4DS-1, del4DS-2, del4DS-3 and del4DS-4, respectively.

### Marker development and genotyping

Homoeologue nonspecific markers were designed to simultaneously amplify fragments of homoeologous genes on 4A, 4B and 4D. Sequence information of 4D genes and corresponding homoeologous genes were obtained from Ensembl Plants (http://plants.ensembl.org/Triticum_aestivum/Info/Index). Gene names and the physical positions reported correspond to the IWGSC RefSeq v1.1 wheat genome assembly (IWGSC, 2018). Sequence insertions and deletions (indels) between homoeologous gene sequences were exploited to enable distinction of the three resulting PCR products. Forward primers were M13-tailed to enable incorporation of a fluorescent adaptor to PCR products, as described by Schuelke (2000). 37 markers designed as such were used to characterise the deletions in four Chinese Spring 4DS terminal deletion lines (Table 2).

**Table 2.**
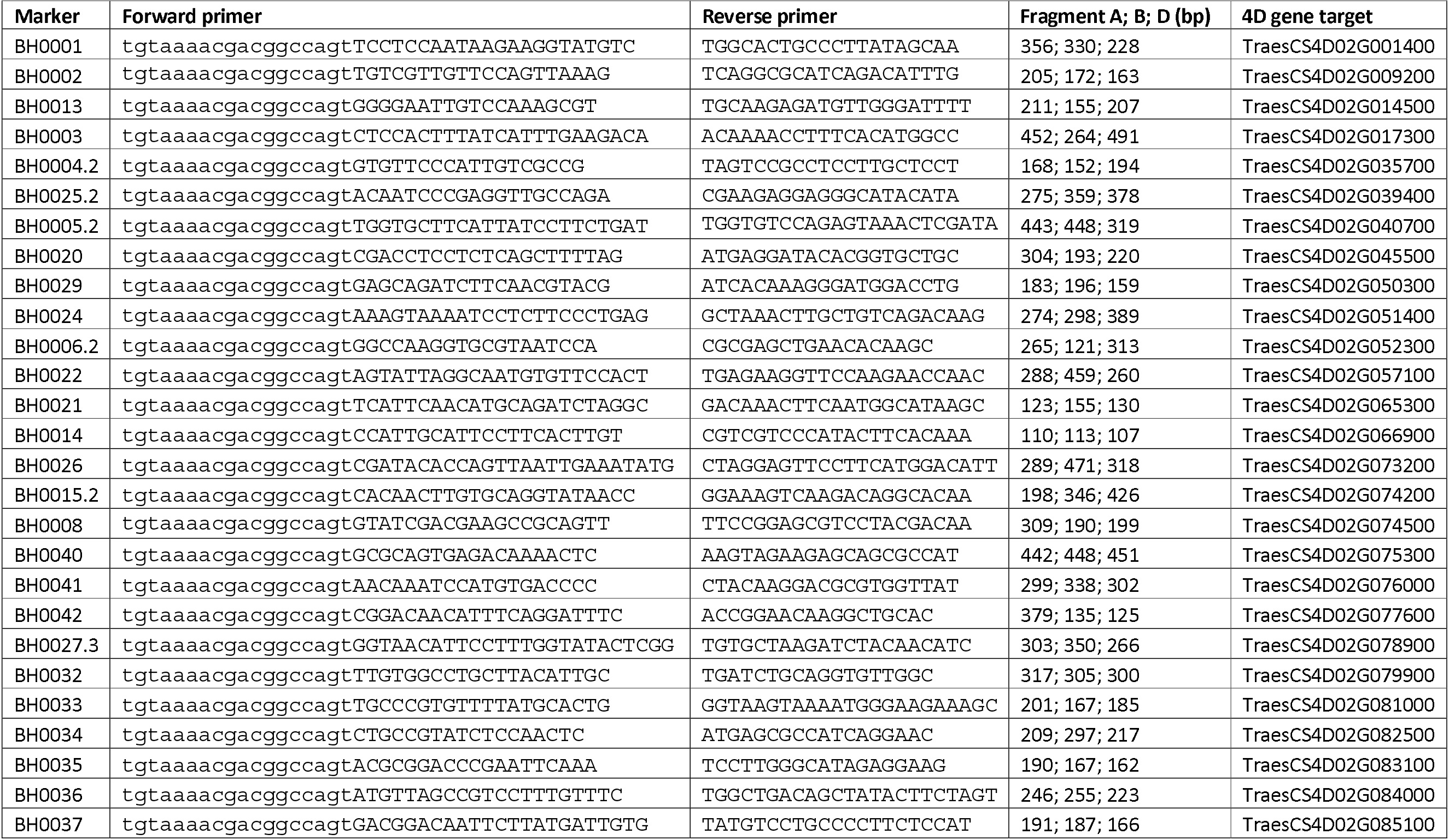
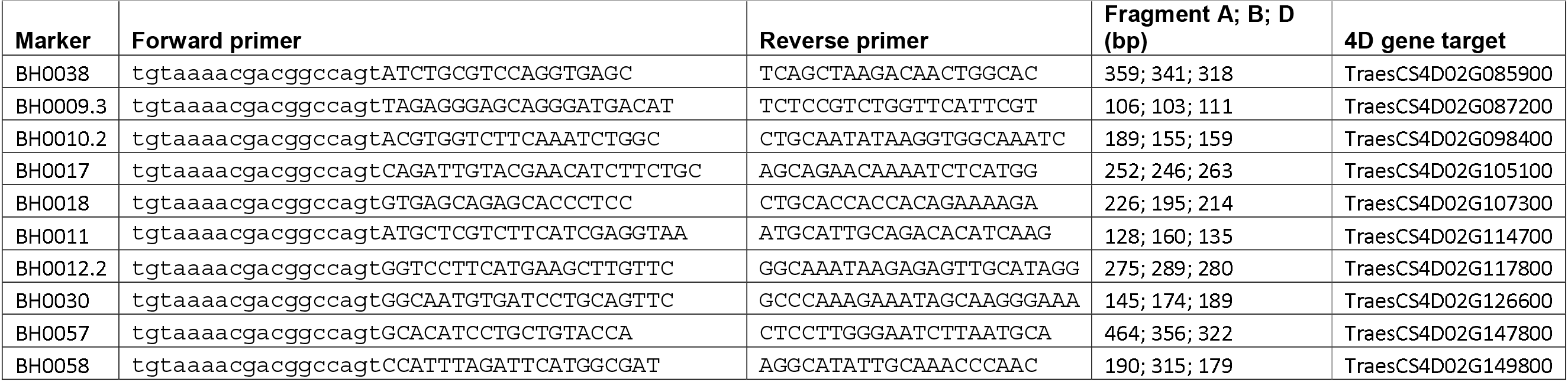
Homoeologue nonspecific markers used to genotype four Chinese Spring 4DS terminal deletion lines. Primer sequences, fragment sizes (corresponding to the 4A, 4B and 4D homoeologous gene targets) and the 4D gene target of markers used to characterise the deletion sizes present in four Chinese Spring 4DS terminal deletion lines. The lowercase sequence in the forward primer indicates the M13 tail. All markers amplified at 58 °C annealing temperature.

DNA was extracted from freeze-dried leaf tissue as described by Pallotta *et al.* (2003). PCR reactions were prepared using HotStarTaq Mastermix (Qiagen) following the manufacturer’s instructions and amplified using the following steps: 95 °C 15 min; 35 cycles of: 95 °C 1 min, 58 °C 1 min, 72 °C 1 min; 72 °C 10 min. PCR products were separated using an ABI 3730xl DNA analyser (Applied Biosystems) and resolved using Peak Scanner 2 software (Applied Biosystems). Up to five markers were multiplexed following PCR to increase assay efficiency.

Primers were designed to specifically amplify within a 5H barley UGT-glucosyltransferase (HORVU5Hr1G047150), whilst avoiding amplification of wheat orthologues (primer sequences: GATGAGGTTTGAGATTTGCGGA, CACGAGCACAACAGATGAATTCA). PCR reactions were prepared using Taq Mastermix (Qiagen) and amplified using the following PCR settings: 94°C 3 min; 35 cycles of: 94 °C 30 sec, 58 °C 30 sec, 72 °C 1 min; 72 °C 10 min. PCR products were separated on a 0.8 % w/v agarose gel.

### FHB evaluation and statistical analysis

Highly virulent DON-producing isolates of *F. graminearum* or *F. culmorum* were used in disease experiments. Production of inoculum was carried out as described previously in Gosman *et al.* (2005). Wheat heads were inoculated at mid-anthesis. The conidial suspension, adjusted to 1 *10^6^ spores ml^-1^, was injected in to a spikelet approximately central on the wheat head. The spread of disease symptoms was scored regularly after inoculation. Polytunnel experiments were organised in a randomised complete block design with four replicates each containing four or five plants per line. For the glasshouse experiment, at least 16 plants per lines were randomised and individual inoculated heads were considered as replicates.

Disease data were analysed using a linear mixed model (REML) in Genstat software (v18.1) to assess the variation attributable to line (fixed), inoculation date (fixed), the interaction between line and inoculation date (fixed), and replicate (random), where factors were significant in the model. Data from which residuals were not normally distributed or where residuals did not appear independent of fitted values were log10 transformed, which was sufficient in correcting for these assumptions. Predicted mean and standard error values were calculated for lines included in the REML. Pairwise comparisons were made between the wild type wheat parent/ genetic background and the other genotypes tested in each experiment using Fisher’s protected least significant difference. All predicted values generated from transformed data were back transformed to the original scale for presentation.

### DON evaluation and statistical analysis

DON was purified to > 98 % at IFA-Tulln, as described by Altpeter and Posselt (1994). DON application was carried out on wheat spikes at mid-anthesis, following a protocol modified from Lemmens *et al.* (2005). Two adjacent spikelets opposite to each other on the wheat head and approximately central on the head, were cut with scissors approximately central on the spikelet. 1-2 h after cutting, 10 μL of DON solution (10 mg / mL amended with 0.01 % v/v Tween 20) was applied to the two outer florets of each cut spikelet, between the palea and lemma. To increase the humidity at the site of DON application, treated wheat heads were bagged. At 48 h post-application, the DON application was repeated, and heads bagged again. Hence, each treated wheat head received a total application of 0.8 mg DON. After a further 48 h, crossing bags were removed from the DON treated heads. The severity of bleaching for each treated wheat head was scored, out of ten, daily between five and nine days post application (from the first application). A score of zero was given when no evidence of DON damage was present and a score of ten was recorded when the spike was completely bleached above the point of DON application. Scores between one and nine were used to record the progressive yellowing and bleaching of the DON treated wheat heads, which occurred relatively uniformly above the point of DON application in the case of Chinese Spring (Figure S1). After the experiment, DON-treated and untreated heads from each plant were harvested. From each plant with a DON treated head, a comparable untreated head (with similar spikelet number and head length) was selected for grain weight analysis. Grain number and grain weight data were collected from DON treated and comparable untreated heads from each plant, to observe any difference in the effect of DON on grain filling.

DON bleaching data and associated grain data were analysed using a REML. Both DON bleaching data and grain data were log10 transformed to achieve normality of residuals and to ensure residuals were independent of fitted values. For bleaching data, line was included as a fixed term and replicate as a random term in the model. For DON grain data, the REML model was constructed using line, treatment (DON treated or untreated heads), and the interaction between line and treatment as fixed terms, and replicate as a random term. Ratios between mean treated and untreated values were calculated by subtracting the predicted mean of log10 DON treated heads from the predicted mean of log10 untreated heads for each line. Standard errors of predicted means were calculated as the square root of the sum of the squared standard errors of the predicted mean values. The calculated mean and standard error values were back transformed, resulting in the presentation of DON treated/ untreated mean grain weight ratios for each line.

## Results

### Effect of barley chromosome additions, substitutions, translocations and centric fusions on type II FHB susceptibility in the winter wheat variety Martonvasari 9 (Mv9kr1)

FHB point inoculation experiments of the wheat-barley material were conducted twice and are described as experiment 1 (Figure 1A) and experiment 2 (Figure 1B) henceforth. The experiments showed very similar results for most of the lines tested. FHB symptoms were always restricted in both barley varieties, Igri and Betzes, and did not spread from the inoculated spikelet. For this reason, Igri and Betzes were only included as control lines in experiment 1 (Figure 1A). The primary wheat parent, Mv9kr1, was susceptible to the spread of the fungus in both repeats of the experiment.

**Figure 1.**
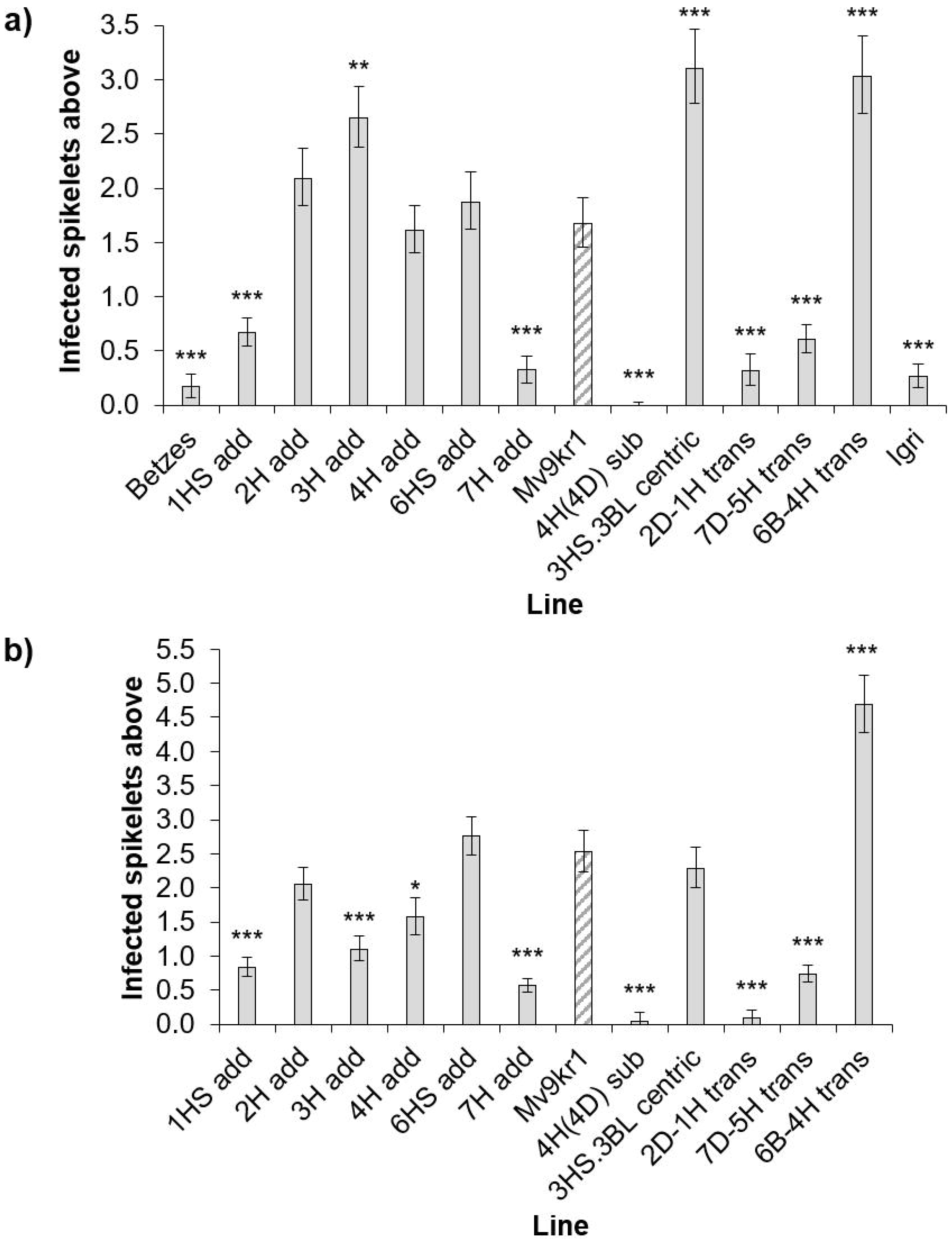
FHB disease above inoculation point in wheat-barley addition, substitution, translocation and centric fusion lines from a) polytunnel experiment 1, including barley parents Igri and Betzes as controls, and b) polytunnel experiment 2. Predicted means were generated using a linear mixed model. Error bars are ± standard error. * p< 0.05; ** p< 0.01; *** p< 0.001 compared to Mv9kr1.

The addition of barley chromosomes 2H (2H add) and 6HS (6HS add) appeared to have no effect on FHB resistance in either experiment. Disease symptoms in these lines were not statistically significantly different from that of Mv9kr1. The 6BS.6BL – 4HL translocation (6B-4H trans) was significantly more susceptible than Mv9kr1 (p< 0.001 in both experiments). Whilst the 3HS.3BL centric fusion line (3HS.3BL centric) was more highly susceptible in experiment 1 (p< 0.001), the line showed similar disease to Mv9kr1 in experiment 2 (p= 0.566). The addition of chromosomes 1HS (1HS add) and 7H (7H add), in addition to the 5HS-7DS.7DL wheat-barley translocation (5H-7D trans) and the 2DS.2DL-1HS translocation line (2D-1H trans) all showed highly significant increases in FHB resistance compared to Mv9kr1 (p< 0.001 in both experiments for all lines). The 3H addition (3H add) was inconsistent between the two experiments. In experiment 1, the 3H addition was significantly more susceptible to FHB than Mv9kr1 (p= 0.004) whilst, in experiment 2, it was significantly more resistant (p< 0.001).

A particularly strong resistant phenotype was seen with the 4H(4D) substitution, in which disease was almost entirely restricted to the inoculated spikelet in both experiments (p< 0.001 in both instances). In contrast to this, the addition of barley 4H (4H add) showed similar disease levels to Mv9kr1 in experiment 1 (p= 0.841, Figure 1A) and exhibited only a small increase in resistance in experiment 2 (p= 0.021, Figure 1B).

### Effect of barley chromosome additions, substitutions, translocations and centric fusions on type II FHB susceptibility in the spring wheat variety Chinese Spring

An FHB point inoculation experiment was performed on wheat-barley addition lines of the varieties Chinese Spring and Betzes, respectively (Figure 2). These lines include addition lines of 5HS and 5HL, which were absent in the lines generated in the Mv9kr1 wheat background. As previously observed, Betzes showed almost no disease spread from the inoculation point. Chinese Spring, on the other hand, showed evidence of disease spread. FHB symptoms in the majority of addition lines were not significantly different from Chinese Spring. The addition lines carrying the barley chromosome arms 2HL, 6HS, 7HL and 7HS all showed significantly increased FHB susceptibility compared to Chinese Spring.

**Figure 2.**
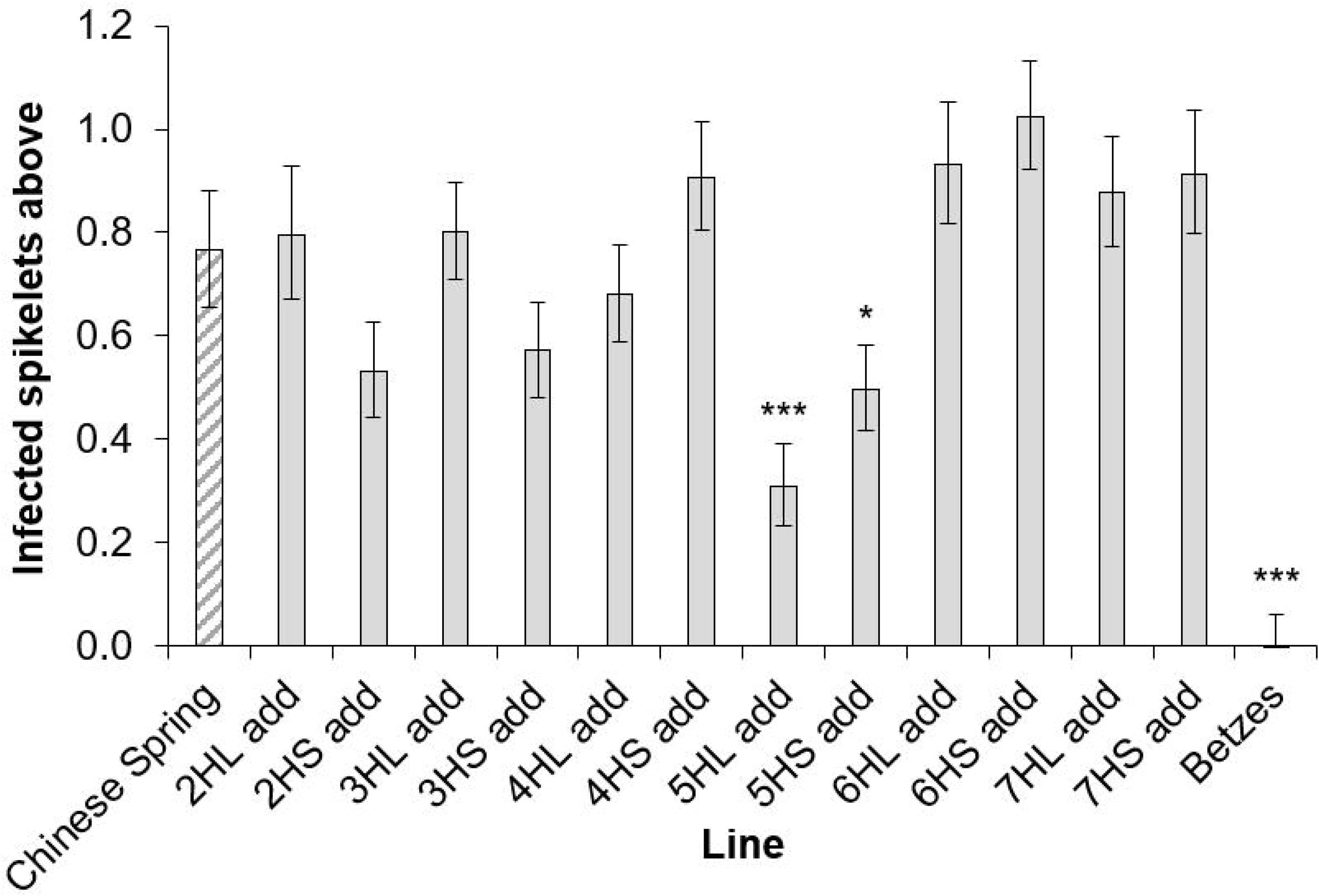
FHB disease, as a percentage of total number of bleached spikelets, from data combined from 13 dpi and 14 dpi. Predicted means were generated using a linear mixed model. Error bars are ± standard error. * p= 0.05-0.01 compared to Chinese Spring; *** p< 0.001 compared to Chinese Spring.

The 5HL addition line exhibited significantly increased FHB resistance when compared with Chinese Spring (p< 0.001), although the line was still significantly more susceptible than Betzes (p= 0.042). The 5HS addition line was also statistically significantly more resistant compared to Chinese Spring (p= 0.039). A marker targeting the barley UDP-glucosyltransferase gene, HORVU5Hr1G047150, confirmed that this gene was present in Betzes and the 5HL addition line, but was absent in the 5HS addition line (Figure S2). Consistent with the previous experiments, the 4HL and 4HS addition lines both showed similar FHB susceptibility to Chinese Spring.

### Type II FHB susceptibility and DON susceptibility in Chinese Spring 4D ditelosomic lines

The contrast in the effect of adding 4H or substituting 4D with 4H indicated that the presence of 4D may be responsible for a significant proportion of the susceptibility of both Mv9kr1 and Chinese Spring. To test this possibility, Chinese Spring and two ditelosomic lines: DT(4DL) and DT(4DS), missing 4DS and 4DL, respectively, were tested in three independent FHB point inoculation experiments. Data is presented here from a 2013 experiment conducted in a glasshouse, but the results were replicated in a 2013 experiment under controlled conditions and in a polytunnel experiment conducted in 2016. Chinese Spring and DT(4DS), missing 4DL, showed very similar disease symptoms to each other (Figure 3). In contrast to this, DT(4DL), missing 4DS, was highly resistant to the spread of infection when compared to wild type Chinese Spring (p< 0.001).

**Figure 3.**
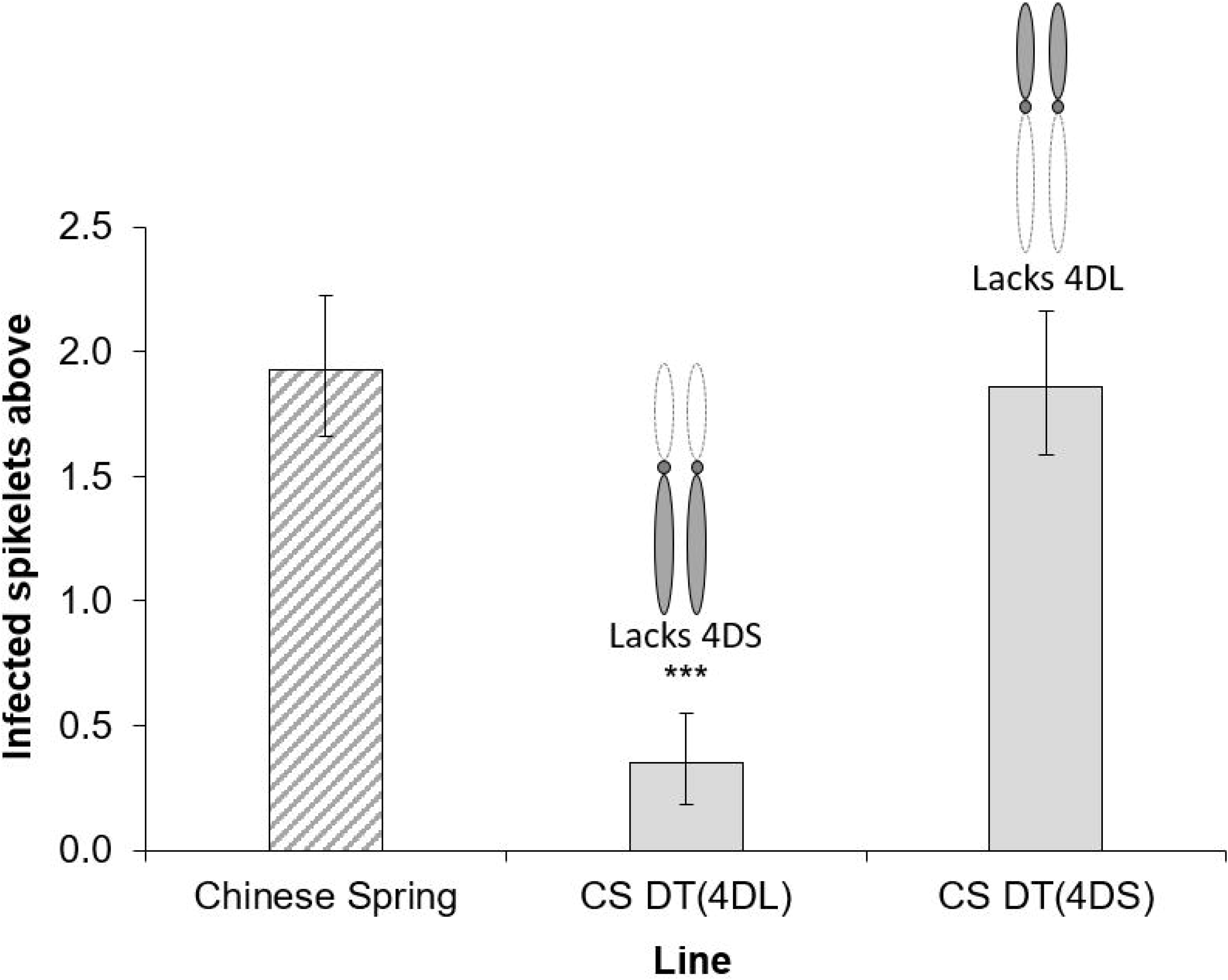
FHB disease at 17 dpi in euploid Chinese Spring and 4D ditelosomic lines DT(4DL) and DT(4DS), missing 4DS and 4DL, respectively. Diagrams of 4D are included above ditelosomic lines to illustrate their genetic state. Error bars are ± standard error. *** p< 0.001 compared to Chinese Spring.

DON is widely believed to contribute towards Type II susceptibility by promoting the spread of FHB. Hence, it is possible that the susceptibility factor may be responding to DON and not the fungus itself. To confirm whether DON is involved, we applied purified DON to wheat heads of Chinese Spring and two ditelosomic lines; DT(4DL) and DT(4DS). Chinese Spring was moderately susceptible to DON, with an average bleaching score of 3.39 (Figure 4A). DT(4DS), lacking 4DL, was not significantly different from Chinese Spring (mean= 2.88; p= 0.222) (Figure 4A). On the other hand, DT(4DL), lacking 4DS, was significantly more susceptible to DON induced bleaching (mean= 7.64; p< 0.001) (Figure 4A).

**Figure 4.**
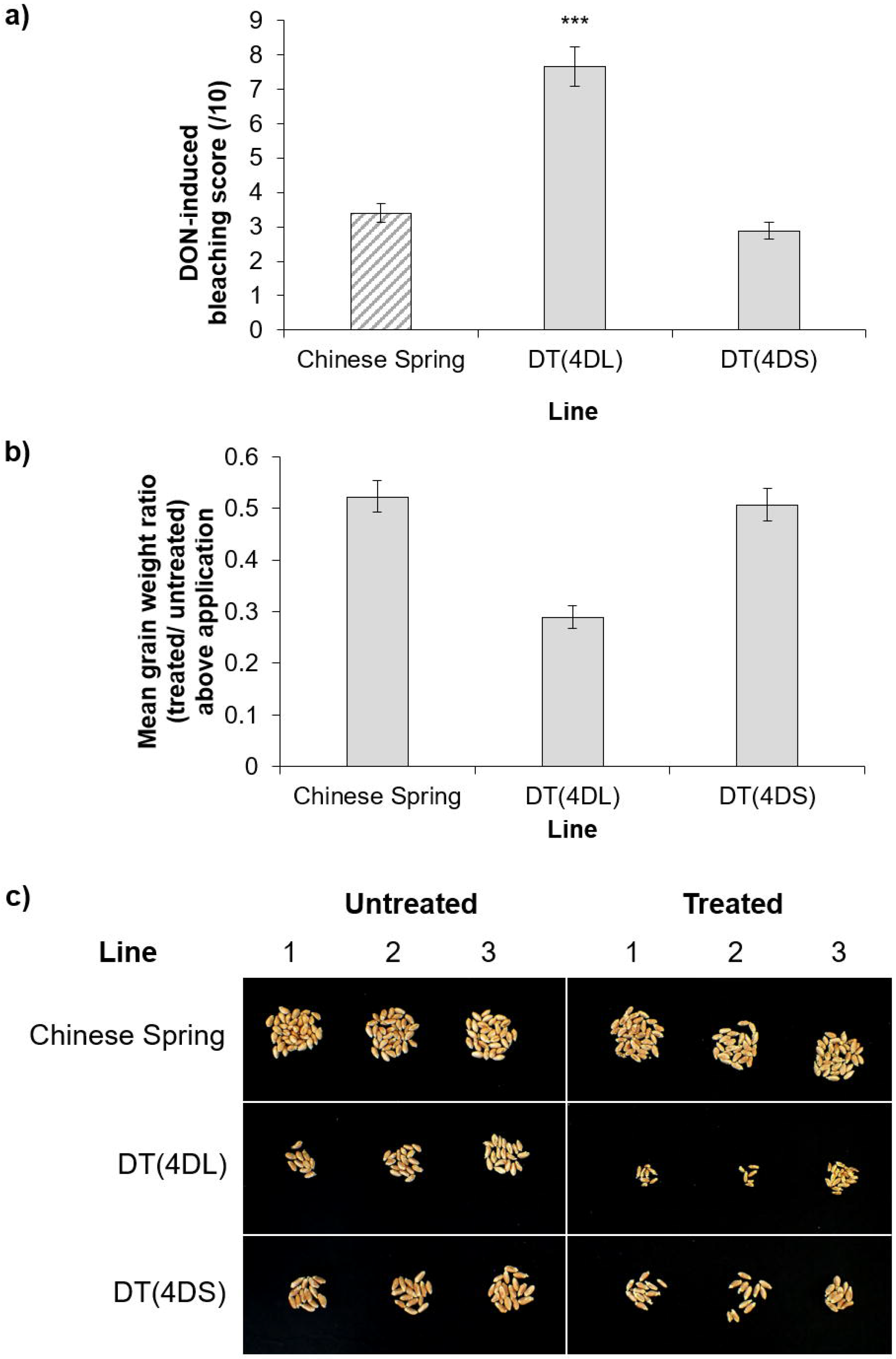
DON application experiment to heads of Chinese Spring and ditelosomic lines DT(4DL) and DT(4DS), lacking 4DS and 4DL, respectively. a) average DON bleaching scores at 7 days post application. Predicted means were generated using a linear mixed model. Error bars are ± standard error. p< 0.001 compared with Chinese Spring. b) ratio of DON treated/ untreated mean grain weight above the DON application point, or comparable point in untreated heads, dissected after the experiment. Ratios were calculated by subtracting the log10 mean grain weight of DON treated heads from untreated heads for each line, followed by back transformation to obtain a treated/untreated ratio for each line. Predicted means were generated using a linear mixed model. Error bars are ± standard error. c) photograph showing three representative examples of untreated and DON treated grain taken from above the DON application point, or comparable point in untreated heads for each line.

Grain was harvested and dissected from DON treated and untreated heads to assess any difference in grain weight. These data closely mirrored the bleaching data. Chinese Spring and DT(4DS) showed similar reductions in grain weight when comparing DON treated and untreated heads (mean ratios of 0.522 and 0.506, respectively) (Figure 4B). In contrast, grain of DON treated DT(4DL) heads had a proportionally much greater reduction in grain weight compared to untreated heads (mean ratio= 0.290) (Figure 4B). The difference is evident when visually comparing treated and untreated grain from the three lines; treated grain from DT(4DL) are visibly smaller than those of Chinese Spring and DT(4DS) (Figure 4C).

These data suggest that DON is not implicated in the function of the susceptibility factor. However, there does appear to be an independent DON resistance factor also on 4DS.

### Precise characterisation of deletion sizes in Chinese Spring 4DS terminal deletion lines

Experiments using 4D ditelosomic lines strongly suggest that the FHB susceptibility attributed to chromosome 4D is isolated to the short arm (4DS). Genotyping was performed on four Chinese Spring lines with terminal deletions on 4DS to verify the deletions present and more precisely position the deletion breakpoint in each line relative to the physical map. Markers were designed that can reliably detect genes on 4D and their homoeologues on 4A and 4B. The ability to detect and distinguish all three homoeologues provides two internal positive controls for each marker when identifying deletions of any particular homoeologue. Up to five markers, tagged using different fluorophores (NED, FAM, PET or VIC), were multiplexed into a single sample for efficiency, using markers designed to produce PCR product sizes sufficiently different for each gene target and its respective homoeologues when resolved using capillary electrophoresis (Figure 5).

**Figure 5.**
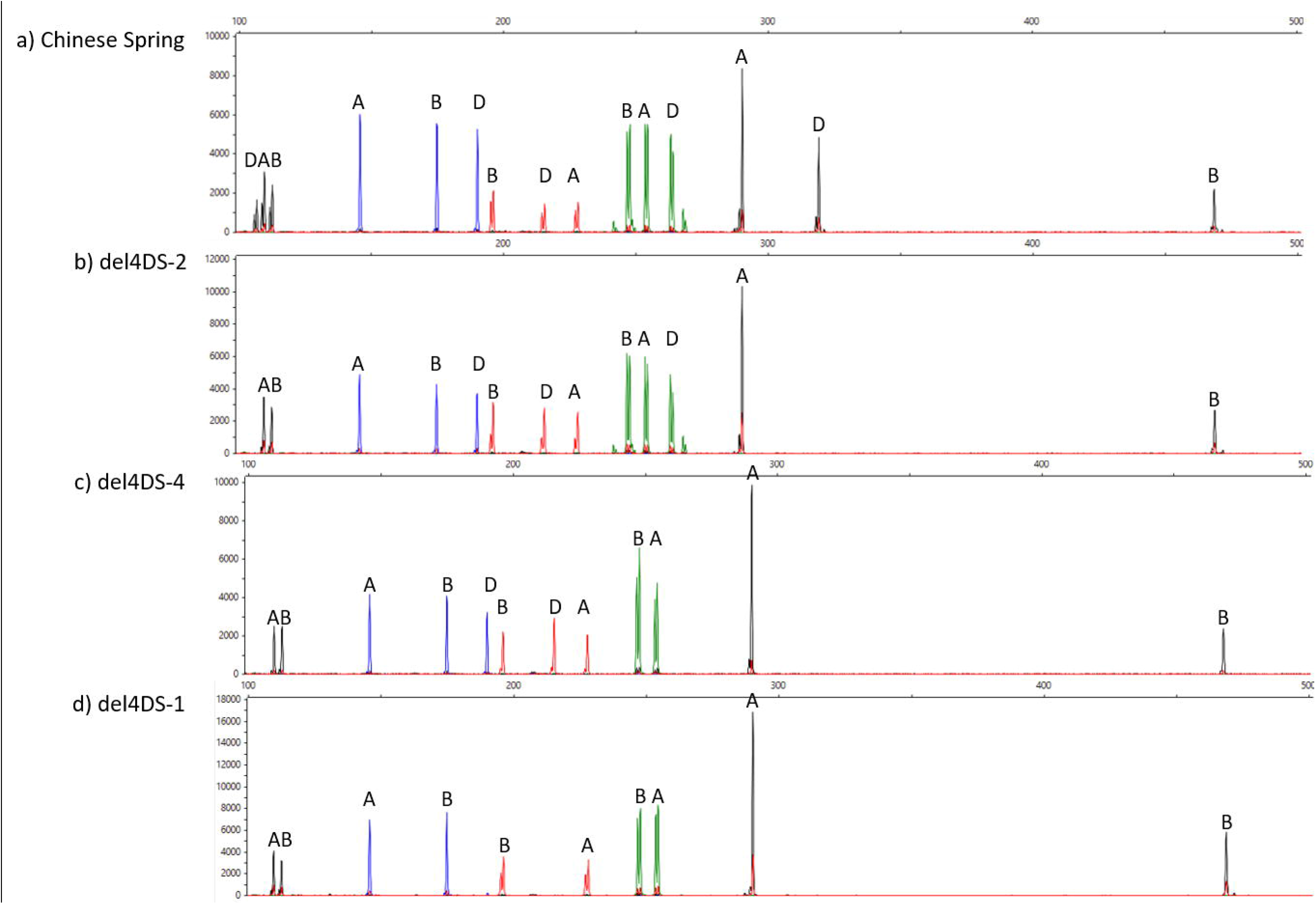
Example outputs of five multiplexed markers BH0014 (left black), BH0030 (blue), BH0018 (red), BH0017 (green) and BH0026 (right black) in a) Chinese Spring; b) del4DS-2; c) del4DS-4; d) del4DS-1. The line del4DS-3 showed the same deletion pattern for the markers visible in the selected multiplex and was hence omitted. X axis is fragment size (bp) and Y axis is the strength of fluorescence (relative fluorescence units). Images were extracted as screenshots from Peak Scanner 2 software (Applied Biosystems).

Genotyping was successful in identifying genes, and their respective physical positions, flanking the deletion breakpoint in all four 4DS terminal deletion lines (Table 3). A marker (BH0001) targeting the gene TraesCS4D02G001400 at the extreme distal end of 4DS confirmed that all four lines were true terminal deletions. The terminal deletion in del4DS-2 extends to between 50.6 and 51.6 Mbp. Line del4DS-4 is deleted up to between 53.9 and 54.8 Mbp. The deletion in del4DS-3 ends between 83.3 and 85.6 Mbp. The deletion breakpoint in the largest terminal deletion line, del4DS-1, ends between 111.1 and 140.9 Mbp.

**Table 3.** Flanking genes and markers of deletion breakpoints in four Chinese Spring 4DS terminal deletion lines. The breakpoint interval is the size of the interval between two adjacent markers where the marker signal was retrieved, indicating the end of the deletion.

| Line | Left flank gene | Left flank marker | Right flank gene | Right flank marker | Breakpoint interval (Kb) |
| --- | --- | --- | --- | --- | --- |
| del4DS-2 | TraesCS4D02G076000 | BH0041 | TraesCS4D02G077600 | BH0042 | 976 |
| del4DS-4 | TraesCS4D02G079900 | BH0032 | TraesCS4D02G081000 | BH0033 | 949 |
| del4DS-3 | TraesCS4D02G105100 | BH0017 | TraesCS4D02G107300 | BH0018 | 2313 |
| del4DS-1 | TraesCS4D02G126600 | BH0030 | TraesCS4D02G147800 | BH0057 | 29776 |

### Chinese Spring 4DS terminal deletion lines and type II FHB susceptibility

Euploid Chinese Spring and the four Chinese Spring 4DS terminal deletion lines genotyped (del4DS-2, del4DS-4, del4DS-3 and del4DS-1, in ascending order of terminal deletion size) were point inoculated in a polytunnel experiment in 2017 (Figure 6). Chinese Spring showed moderate levels of disease in this experiment, with mean disease above the inoculation point of 1.84 bleached spikelets at 13 dpi. Lines del4DS-2 (p= 0.796) and del4DS-4 (p= 0.278) showed similar disease levels to that of euploid Chinese Spring (Figure 6 and Figure 7). Lines del4DS-3 and del4DS-1 both had significantly reduced disease with respect to euploid Chinese Spring (p< 0.001 for both lines) (Figure 6 and Figure 7).

**Figure 6.**
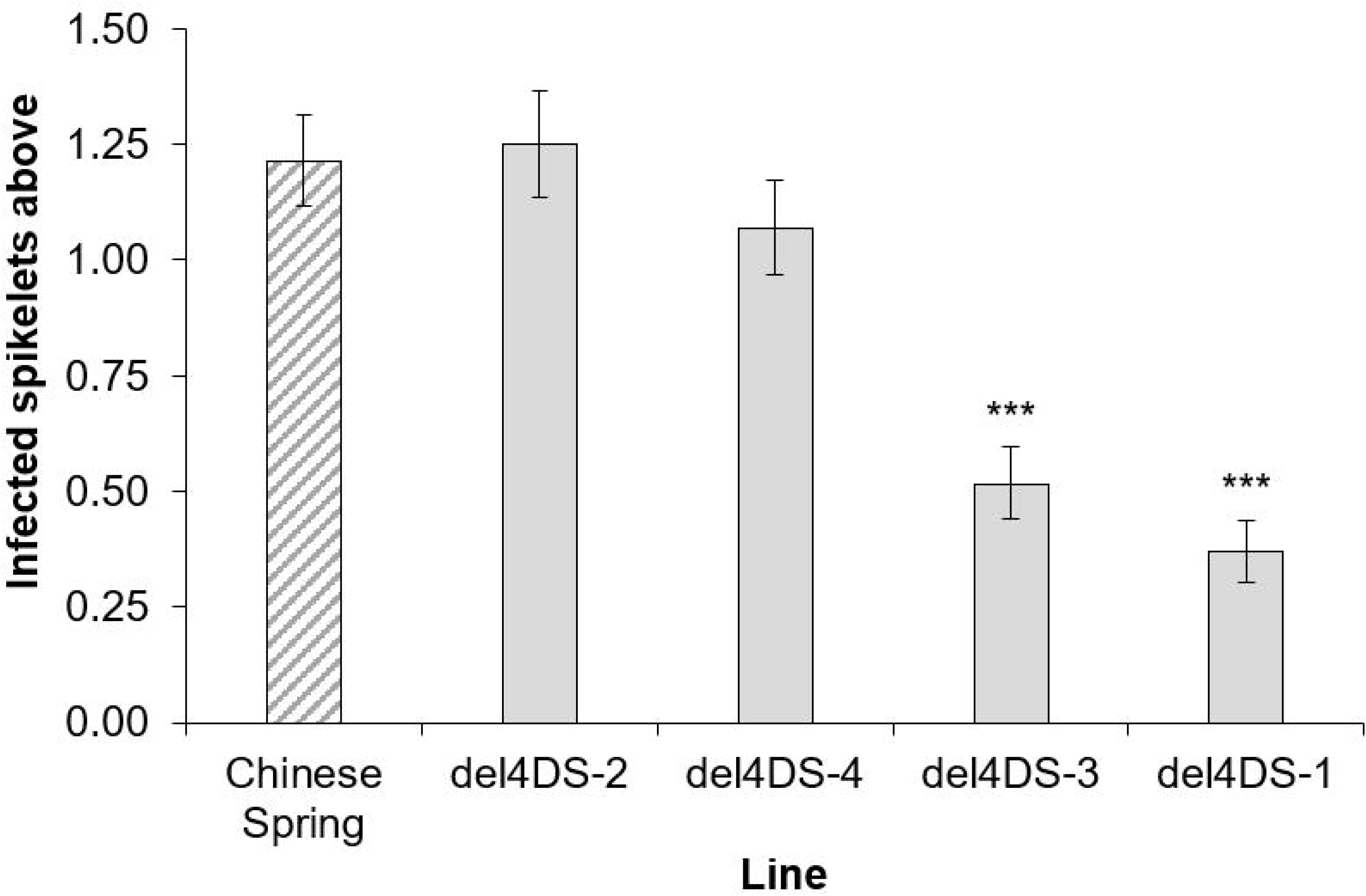
FHB disease above the inoculation point at 13 dpi, following point inoculation of euploid Chinese Spring and four terminal deletion bins; del4DS-2, del4DS-4, del4DS-3 and del4DS-1. Error bars are ± standard error. *** p< 0.001 compared to Chinese Spring.

**Figure 7.**
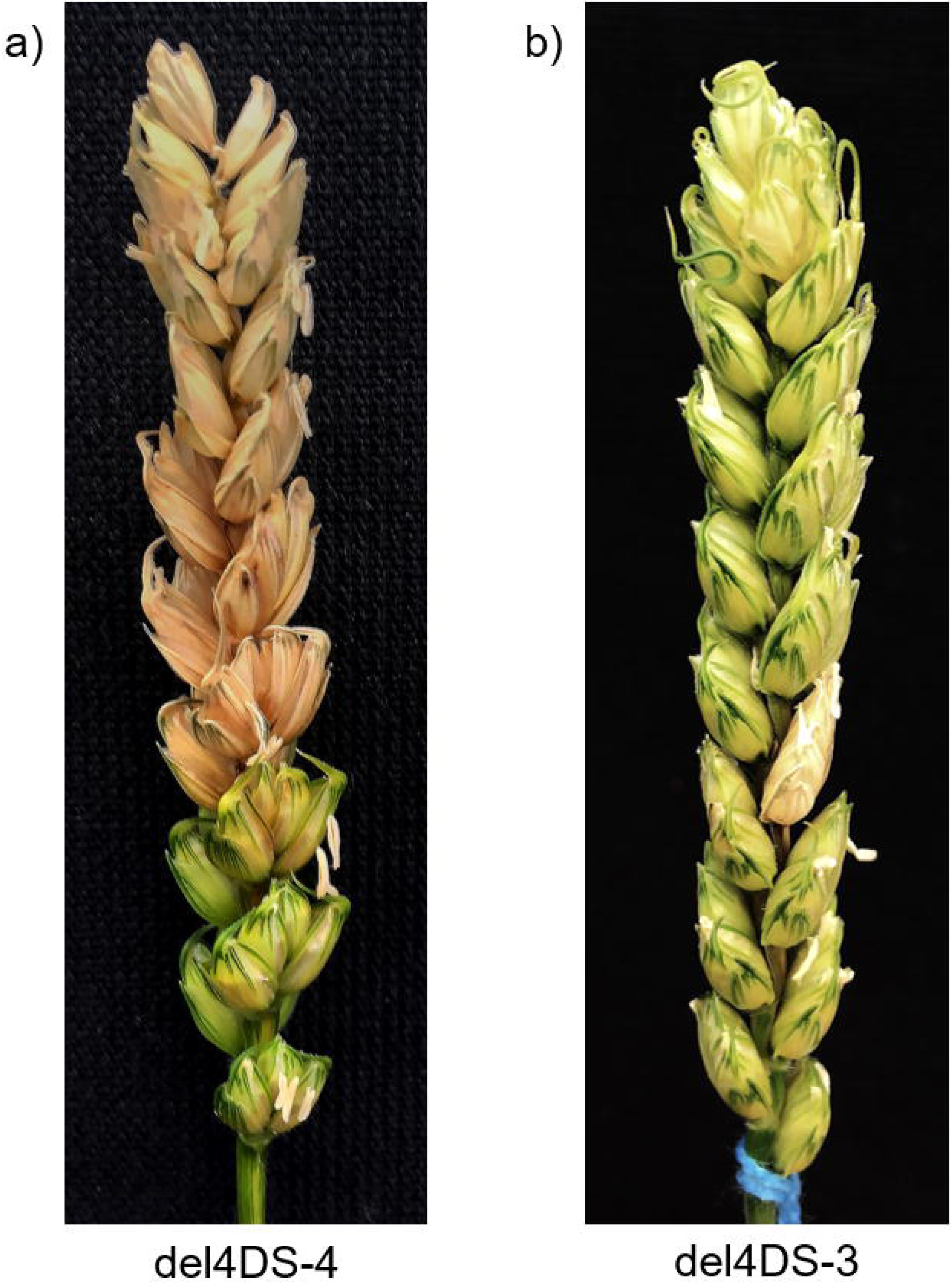
Representative FHB disease symptoms in the Chinese Spring terminal deletion lines del4DS-4 and del4DS-3 at 16 dpi.

This information was used to infer that the susceptibility factor was present in the two deletion lines carrying the smaller deletions (del4DS-2 and del4DS-4) but was lost in the two lines containing the larger deletions (del4DS-3 and del4DS-1). Hence, the FHB susceptibility factor appears to reside between the deletion breakpoints of del4DS-4 and del4DS-3; a 31.73 Mbp interval (Figure 8).

**Figure 8.**
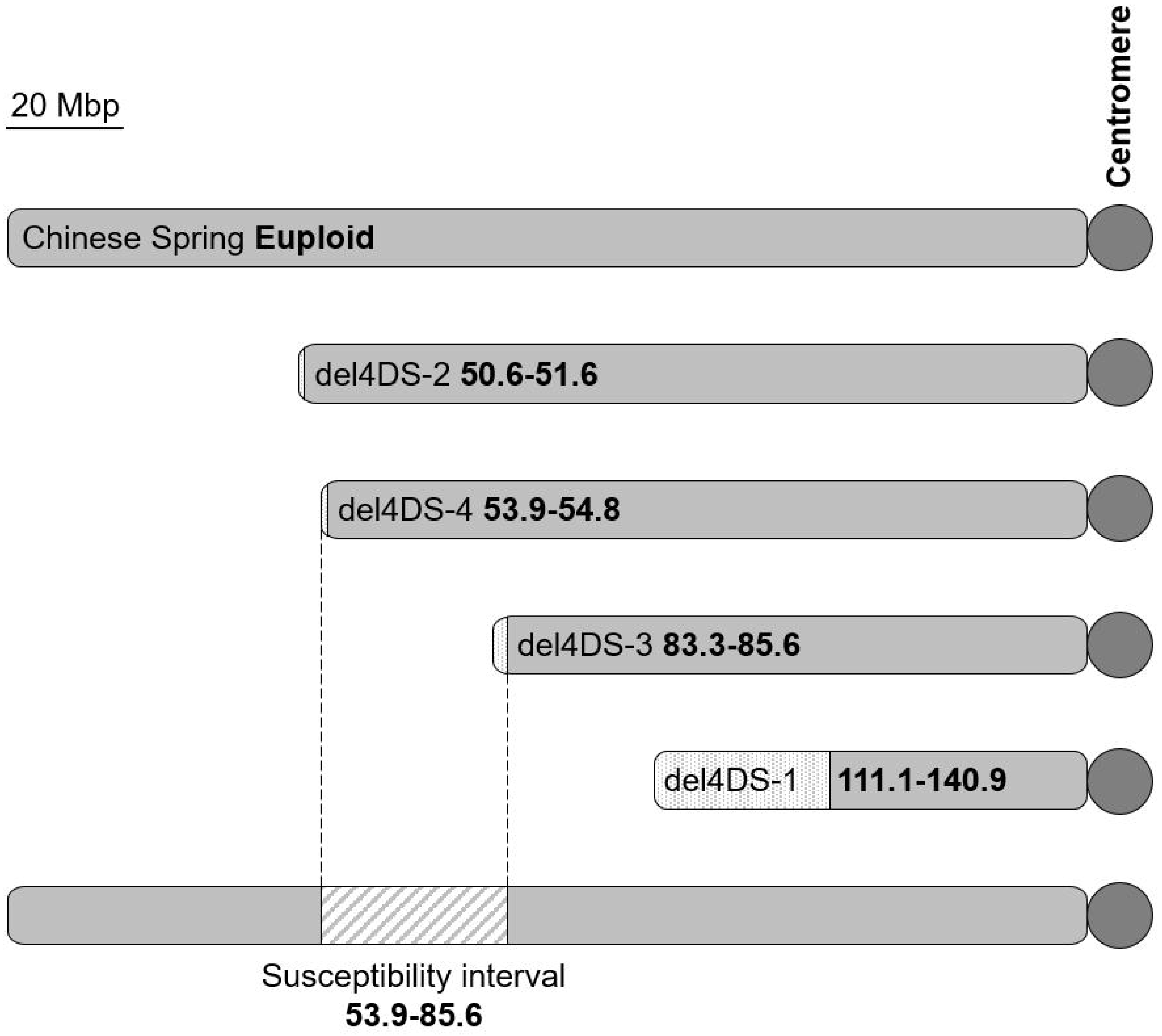
Diagrams of 4DS in euploid Chinese Spring and four 4DS terminal deletion lines, as characterised by genotyping with 35 markers spanning 4DS. The spotted interval indicates the breakpoint interval; the distance between two markers where the 4D signal was retrieved. The bottom diagram indicates the interval on 4DS inferred to contain an FHB susceptibility factor (diagonal stripes), following point inoculation of the Chinese Spring terminal deletion lines. Values in bold indicate the physical position in Mbp.

## Discussion

Previous studies have shown that barley is able to detoxify DON through glucosylation by the UDP-glucosyltransferase UGT13248 (Schweiger *et al.*, 2010). This gene has been transgenically expressed in Arabidopsis where it was demonstrated to increase resistance to DON (Schweiger *et al.*, 2010). Furthermore, expression of UGT13248 in wheat, under the maize ubiquitin promoter, increased FHB resistance and transformants were demonstrated to more efficiently convert DON to the less toxic DON-3-O-glucoside (Li *et al.*, 2015). However, Xing *et al.* (2018) demonstrated that overexpression of a wheat UGT-glucosyltransferase also increased FHB resistance and reduced the DON concentration in grain. How the barley UDP-glucosyltransferase performs in wheat under its native barley promoter has not yet been demonstrated and hence the increase in resistance attributed to the barley UGT-glucosyltransferase in wheat may be due to overexpression. The barley UDP-glucosyltransferase UGT13248 is encoded by gene HORVU5Hr1G047150 which is present near the centromere on chromosome 5H (Ensembl Plants). If the breakpoints in the wheat - barley 5HS and 5HL ditelosomic addition lines are not centromeric, this may explain the findings related to the high level of resistance conferred by addition of both 5HS and 5HL. To confirm this, we designed primers specific to the barley copy of the UDP-glucosyltransferase and will not amplify from the orthologous wheat copies in the wheat-barley additions. This assay confirmed that the UDP-glucosyltransferase was isolated to the 5HL addition line and was absent in the 5HS addition line. Hence, it is likely that an independent source of FHB resistance is present on 5HS.

In this study, we also found that addition of the barley chromosome 7H (7H add) or the short arm of chromosome 1H (1HS add), as well as the translocation of 1H to 2D (2D-1H trans), significantly increased Type II FHB resistance in the winter wheat variety Mv9kr1. Despite the enhanced FHB resistance from the addition of 7H to Mv9kr1, the addition of neither 7HS nor 7HL had an effect in the Asian spring wheat cultivar Chinese Spring. No 1H addition lines were available in the Chinese Spring-Betzes addition set, so this could not be compared between populations. These findings suggest that barley contains genes conferring Type II resistance that are lacking in one or both wheat varieties. The addition of barley chromosomes 5H and perhaps 1H and 7H are likely to offer the best opportunity of enhancing FHB resistance, when considering the use of wheat-barley introgressions.

We confirmed the presence of a possible Type II susceptibility factor on the short arm of 4D in three independent experiments. The loss of 4DS (line DT(4DL)) resulted in a high level of FHB resistance, whilst the loss of 4DL (line DT(4DS)) resulted in little change compared to euploid Chinese Spring. Ma *et al.* (2006) phenotyped Chinese Spring ditelosomic lines for FHB susceptibility and they also reported an increase in FHB resistance in the line missing 4DS. Together, these studies strongly suggest the presence of a susceptibility factor in both winter (Mv9kr1) and spring (Chinese Spring) wheat genetic backgrounds. We applied purified DON to the 4D ditelosomic lines to test whether or not the susceptibility factor is being influenced by DON. However, the loss of 4DS resulted in higher susceptibility to DON, assessed both by scoring DON induced bleaching and by comparing grain weights. This would indicate that there is an independent resistance factor to DON present on 4DS and that the susceptibility factor is increasing susceptibility to the fungus or another virulence factor.

Endo and Gill (1996) developed a set of terminal deletion lines in Chinese Spring. The lines have deletions from the ends of each chromosome arm, varying in size. These stocks are a valuable resource for physically mapping genes to a defined interval of a chromosome arm. The lines were characterised using C-banding and the deletion size reported as a fraction length (FL) value; effectively the proportion of the chromosome arm estimated to have been retained. C-banding is unlikely to be capable of reliably detecting more complex deletions, such as interstitial deletions or chromosome substitutions. Since their development, the Chinese Spring terminal deletion stocks have not been more precisely characterised using more recent advancements in genotyping. We have genotyped four lines containing terminal deletions of 4DS, using a total of 37 novel homoeologue nonspecific markers spanning the chromosome arm. These markers take advantage of the hexaploid nature of wheat to create a robust genotyping assay for the detection of deletions on 4DS, and its homoeologous regions on 4BS and 4AL. A similar assay was used by Chia *et al.* (2017) to verify deletions across homoeologous regions but this study expands on this technique, using a much higher density of markers to characterise deletion size. Homoeologous genes are simultaneously amplified with a single pair of primers but are distinguishable due to differences in the size of PCR products corresponding to the A, B and D genome copies. The signal from the retained homoeologues act as internal controls for a deletion in any homoeologue; in this case, the 4D copy. This technique verified that all four Chinese Spring 4DS terminal deletion lines were indeed true terminal deletions and the size of the deletions were consistent with the FL values calculated by Endo and Gill (1996). For the lines del4DS-2, del4DS-4 and del4DS-3, the physical position of the deletion endpoint has been restricted to a small interval. For both del4DS-2 and del4DS-4, this interval is smaller than 1 Mbp. The interval containing the deletion endpoint in del4DS-3 has been refined to approximately 2.3 Mbp. The breakpoint in the largest deletion, del4DS-1, was less precisely characterised and the deletion breakpoint was isolated to a 29.8 Mbp interval. For the purposes of this study, it was not necessary to more precisely characterise the deletion in del4DS-1, because the FHB susceptibility factor appears to be situated between the deletion breakpoints in lines del4DS-4 and del4DS-3.

We performed FHB disease experiments on the four Chinese Spring 4DS terminal deletion lines that we genotyped. This clearly demonstrated that the lines with the two smaller deletions, del4DS-2 and del4DS-4, retained the susceptibility factor and showed a similar phenotype to euploid Chinese Spring. In contrast the lines del4DS-3 and del4DS-1, containing the larger deletions, showed significantly improved FHB resistance and hence the susceptibility factor has presumably been lost. As the susceptibility factor was present in del4DS-4 but was lost in del4DS-3, it must be situated between the deletion breakpoints of these two lines, restricting the susceptibility factor to a 31.7 Mbp interval containing 274 high confidence genes (IWGSC RefSeq v1.1). The positive effect of the deletion of the susceptibility factor appears to be restricted to 4D and hence it is likely the gene responsible is 4D specific and does not possess homoeologues. BLAST searches of each 4D gene in the interval identified 20 genes that appear to lack homoeologues and hence are 4D-specific. Alternatively, the 4D homoeologue may be preferentially expressed compared to the 4A and 4B copies. It is also possible that the improved FHB resistance is the consequence of altered dosage of the 4D susceptibility factor and its homoeologues. The disrupted balance of a physiological process exploited by the fungus is also likely to result in altered disease susceptibility.

A population possessing smaller deletions is required to further refine the position of the FHB susceptibility factor. We intend to utilise a gamma-irradiated population of the UK spring wheat variety Paragon (Shaw *et al.*, 2013; Wheat Genetic Improvement Network, 2019) to improve the resolution for the physical mapping of the FHB susceptibility factor.

It may be considered surprising that an FHB susceptibility factor with such a powerful effect has not been detected before now. However, we hypothesise that the FHB susceptibility factor is highly conserved among wheat cultivars. The susceptibility factor exists both in the Hungarian winter wheat cultivar Martonvasari 9 and in the Asian spring wheat variety Chinese Spring. Preliminary experiments of gamma irradiated Paragon lines, containing a deletion of the entire 31.7 Mbp FHB susceptibility interval, indicated this line possesses potent resistance and hence confirms that the susceptibility factor is also present in the UK spring cultivar Paragon (data not shown). If there was sufficient allelic variation at the locus, the effect of the susceptibility factor is likely to have been detected as an FHB QTL in existing mapping populations. In the absence of such reports, we predict that the FHB susceptibility factor is fixed in both spring and winter wheats.

Genetic resistance to fungal diseases is critical to the protection of food crops such as wheat. The search and incorporation of resistance factors is common practice in crop plant breeding. However, identifying novel sources of resistance to FHB is challenging and time consuming. FHB resistance is quantitative, highly polygenic, and often environmentally labile. Few large effect FHB QTL have been identified. Attempts to clone the gene underlying the best known source of FHB resistance, the *Fhb1* QTL, have been inconsistent and controversial (Ma *et al.*, 2017; Rawat *et al.*, 2017; Steiner *et al.*, 2017; Su *et al.*, 2017). Rawat *et al.* (2016) reported that they had cloned a pore-forming toxin-like (*PFT*) gene underlying the *Fhb1* QTL. However, Jia *et al.* (2018) disputed the findings of Rawat *et al.* (2016). Su *et al.* (2018) identified that the presence of a deletion at the 5’ end of a histidine-rich calcium-binding protein within the *Fhb1* locus was sufficient in identifying varieties carrying *Fhb1.* Su *et al.* (2019) have since reported that *Fhb1* possesses enhanced resistance due to the loss-of-function of the histidine-rich calcium-binding protein and the wild type allele is hence functioning as a susceptibility factor. Li *et al.* (2019) also identified that mutation of the histidine-rich calcium-binding protein as the gene responsible for *Fhb1* resistance. However, in conflict with the findings of Su *et al.* (2019), their data suggests that this is due to a gain-of-function resulting from an different start codon positioned upstream to the original (Li *et al.*, 2019). Our data on the 3HS-3BL centric fusion line does not suggest that 3BS contains a susceptibility factor, as the line was either wild type-like or more highly susceptible to the spread of FHB. Furthermore, Ma *et al.* (2006) reported that the Chinese Spring ditelosomic line missing 3BS (DT(3BL)) was more susceptible to FHB, which is not compatible with the hypothesis that FHB resistance from *Fhb1* being a loss-of-function susceptibility factor. It remains possible that more than one gene is responsible for FHB resistance conferred by *Fhb1.* Furthermore, it has proven difficult to utilise *Fhb1* in elite varieties in high yielding European environments with few varieties released containing the resistance. This suggests a linkage drag from the resistance or a pleiotropic effect and demonstrates a need for novel methods of conferring resistance, such as eliminating susceptibility factors.

Despite this, there has been relatively little research into susceptibility factors in wheat and other cereals and how they may be used in plant breeding. The barley *mildew resistance locus o (Mlo*) is one of the earliest and best characterised examples of how disruption of a susceptibility factor could be exploited to improve disease resistance; in this case, to powdery mildew caused by the biotrophic fungus *Blumeria graminis* f. sp. *hordei* (Jorgensen, 1992). Induced and natural mutation of the *Mlo* locus result is a recessive, race nonspecific and durable resistance which has been widely deployed in European spring barley varieties (Jorgensen, 1992; Lyngkjaer and Carver, 2000; McGrann *et al.*, 2014). Mlo-based resistance has since been demonstrated in a number of other species affected by powdery mildew, reviewed by Kusch and Panstruga (2017). The deployment of *mlo* in wheat is more challenging due to its allohexaploid nature (Acevedo-Garcia *et al.*, 2017). However, TALENs and CRISPR Cas9-derived gene knockouts (Wang *et al.*, 2014) and *Mlo* TILLING mutants (Acevedo-Garcia *et al.*, 2017) have been used to demonstrate that mutation of all wheat copies strongly enhances resistance to wheat powdery mildew.

R genes, usually nucleotide binding site-leucine rich repeat (NBS-LRR) genes, are typically used by plants to detect and respond to attack by biotrophic fungi. However, necrotrophic pathogens have evolved methods of exploiting such plant defences to aid infection. *Parastagonospora nodorum* and *Pyrenophora tritici-repentis* are necrotrophic pathogens of wheat that utilise this strategy. Susceptibility to these diseases operates in an inverse gene-for-gene interaction, in which a fungal necrotrophic effector is detected by a corresponding host sensitivity gene product (usually an NBS-LRR), triggering a hypersensitive response that results in necrosis that benefits the fungus (Faris *et al.*, 2010). If either necrotrophic effector or host sensitivity gene is absent, the interaction is impossible and host resistance is maintained. There have been few reports of how NBS-LRRs are involved in interactions with *Fusarium* spp. However, Zhang *et al.* (2019) found that the expression of an LRR gene appeared to increase susceptibility to *F. graminearum* in soybean (*Glycine max).*

*Fusarium graminearum* leads a hemibiotrophic lifestyle whereby the hyphal front remains surrounded by living tissue but cell death is triggered soon after colonisation (Brown *et al.*, 2010). Phytohormones play important roles in defence and there is considerable evidence indicating that *F. graminearum* modifies phytohormone expression for its own benefit. Disruption of ethylene signalling in wheat (Chen *et al.*, 2009) and brassinosteroid signalling in barley and *Brachypodium distachyon* (Goddard *et al.*, 2014) results in enhanced resistance to FHB infection, suggesting that the fungus is exploiting phytohormone signalling in order to aid infection. Expression of 9-lipogenases are also manipulated by *F. graminearum* in both bread wheat and *Arabidopsis thaliana* and are hence operating as susceptibility factors (Nalam *et al.*, 2015).

In this study, we provide compelling evidence for the presence of an FHB susceptibility factor on the short arm of chromosome 4D. We have demonstrated that the removal of the susceptibility factor is sufficient to significantly improve Type II FHB resistance and have refined its position to a 31.7 Mbp interval containing 274 high confidence genes. We have designed markers that can reliably detect deletions on 4DS. A subset of these markers covering the susceptibility interval will be utilised in further studies to identify lines containing relatively smaller deletions across the FHB susceptibility interval in a gamma irradiated Paragon population. This will reduce the number of gene candidates for the FHB susceptibility and may lead to the identification of the causal gene.

## Acknowledgements

The authors would like to thank BBSRC (grant number: BB/M016919/1) and RAGT Seeds for supporting the PhD studentship of BH. This work was supported by BBSRC and the John Innes Foundation.

## Abbreviations

dpi: Days post inoculation
DON: Deoxynivalenol
D3G: DON-3-O-glucoside
DT: Ditelosomic
FHB: Fusarium head blight
FL: Fraction length
QTL: Quantitative trait locus
UGT: UDP-glucosyltransferase

